# Cardiolipin, and not monolysocardiolipin, preferentially binds to the interface of Complexes III and IV

**DOI:** 10.1101/2022.07.21.500943

**Authors:** Robin A. Corey, Noah Harrison, Phillip J. Stansfeld, Mark S.P. Sansom, Anna Duncan

## Abstract

The mitochondrial electron transport chain comprises a series of protein complexes embedded in the inner mitochondrial membrane that generate a proton motive force via oxidative phosphorylation, ultimately generating ATP. These protein complexes can oligomerize to form larger structures called supercomplexes. Cardiolipin (CL), a conical lipid, unique within eukaryotes to the inner mitochondrial membrane, has proven essential in maintaining the stability and function of supercomplexes. Monolysocardiolipin (MLCL) is a CL variant that accumulates in people with Barth syndrome (BTHS). BTHS is caused by defects in CL biosynthesis and characterised by abnormal mitochondrial bioenergetics and destabilised supercomplexes. However, the mechanisms by which MLCL causes pathogenesis remain unclear. Here, multiscale molecular dynamics characterise the interactions of CL and MLCL with yeast and mammalian mitochondrial supercomplexes containing Complex III (CIII) and Complex IV (CIV). Coarse-grained simulations reveal that both CL and MLCL bind to sites at the interface between CIII and CIV of the supercomplex. Free energy perturbation calculations show that MLCL interaction is weaker than that of CL and suggest that interaction with CIV drives this difference. Atomistic contact analyses show that, although interaction with CIII is similar for CL and MLCL, CIV makes more contacts with CL than MLCL, demonstrating that CL is a more successful “glue” between the two complexes. Simulations of the human CIII2CIV supercomplex show that this interface site is maintained between species. Our study suggests that MLCL accumulation in people with BTHS disrupts supercomplex stability by formation of relatively weak interactions at the interface lipid binding site.

## Introduction

The mitochondrial electron transport chain is responsible for the generation of ATP at the mitochondrial inner membrane. It includes the respiratory complexes cytochrome *bc*_1_ (or complex III, CIII) and cytochrome *c* oxidase (or complex IV, CIV). CIII catalyses the oxidation of the lipid soluble co-factor ubiquinol and simultaneously reduces the mobile electron transporter cytochrome *c* (cyt *c*). CIV transfers electrons from cyt *c* to oxygen to form water^1^. Both complexes can co-assemble into larger supercomplex structures, although the composition and stoichiometry varies^2^. While the functional significance of supercomplexes remains unsettled, many hypotheses have been proposed. It has been suggested that formation of supercomplexes may increase the stability of individual respiratory complexes; reduce production of harmful reactive oxygen species; or increase electron transfer efficiency between complexes^2^. Recently, it has been proposed that the association of CIII and CIV into a supercomplex structure reduces the distance that the mobile electron carrier, cyt *c*, has to travel between the two^3,4^. Another explanation for supercomplex existence postulates that the structures allow for the unusually high concentration of proteins seen in the inner mitochondrial membrane (IMM). Supercomplex formation is therefore thought to reduce the energetic costs of protein crowding and prevent unproductive protein aggregation^5,6^.

Cardiolipin (CL) constitutes up to 20% of the IMM phospholipid content^7^ and exhibits unique molecular properties as a result of its small headgroup, consisting of two phosphate groups linked by a glycerol bridge, bound to four acyl chains. Direct protein-CL interactions have been shown to influence both protein stability and activity in a range of mitochondrial processes, including generation of the proton motive force as well as apoptosis, mitophagy and mitochondrial fission^8^. CL has been identified in structures of both CIII^9,10^ and CIV^11–13^, and has been shown to be required for their full activity^10,14–19^. CL has further been suggested to act as a ‘glue’ between respiratory complexes, stabilising and promoting association of the individual complexes into the so-called supercomplex structures^20–23^.

Supercomplex structures vary in their composition, often comprising a CIII dimer, one or two copies of CIV and, in the case of the so-called ‘respirasome’, also including Complex I (CI; NADH dehydrogenase). *S. cerevisiae* lacks CI^24^, thus the recently resolved CIII_2_CIV_2_ supercomplexes^3,25–27^ represent a maximal supercomplex. The CIII:CIV interface differs in these supercomplexes, as compared to those containing CI^2,28^, as is also the case in the first mammalian CIII_2_CIV_1_ structures^29^. In the majority of the CIII_2_CIV_1-2_ structures, bound CL is located at the interface between individual complexes^3,26,27,29^ and is able to form polar contacts between both CIII and CIV (Figure 1A).

**Figure 1.**
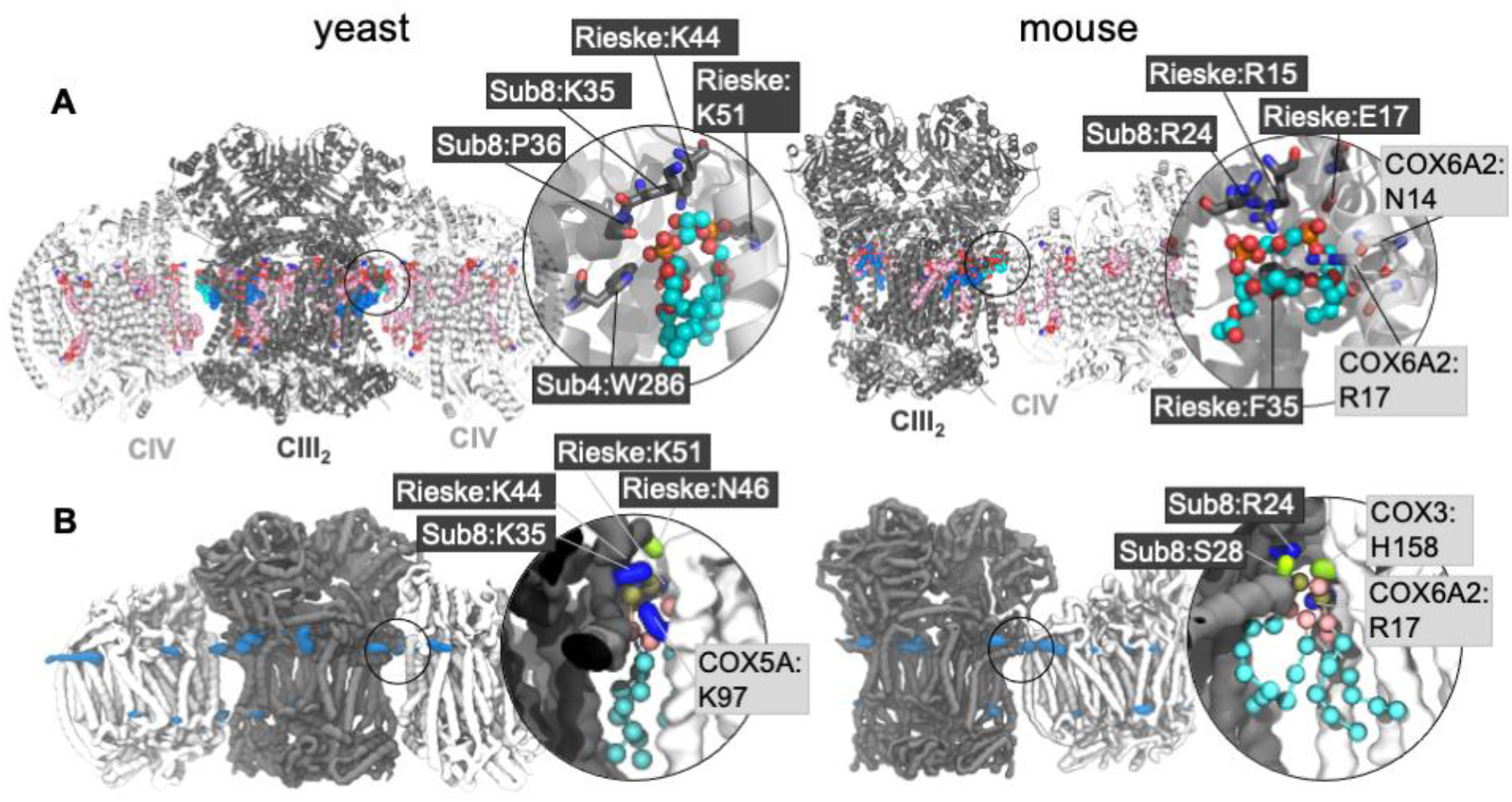
**A)** EM structures of yeast (PDB 6HU9) and mouse (7O3C) supercomplexes, with CL and other bound lipids shown. Non-interface CL are shown in dark blue with the interface CDL is in cyan; other lipids are in pink. Inset is a close-up of the interface CL, with key coordinating residues shown (for comparison with the CG simulations this is defined as residues within 5 Å of the phosphorous atoms of the phosphate moieties and the C1 atom of the bridging glycerol moieties). Phosphorous atoms are in orange; oxygen in red. Residues are labelled with their subunit and name, with black boxed residues from CIII, and pale grey boxed residues from CIV. **B)** CL headgroup density from CIII2CIV2 CG MD simulations of yeast and murine supercomplexes. Insets show snapshots of CL in the interface interaction site. The backbone is shown of CIII (grey) and CIV (white), with sidechains shown of residues within 5.2 Å of the CL headgroup (the phosphate beads and bridging glycerol); lysine and arginine sidechains in blue, histidine, serine and asparagine in green, and phenylalanine in orange. CL is shown with phosphate moieties in tan, glycerol moieties in pink, and acyl chain beads in cyan.

Previous coarse-grained (CG) MD simulations investigating the CL-dependent assembly of CIII_2_CIV_1-2_ supercomplexes show binding of CL at defined binding sites prior to oligomerisation. Following enrichment of CL on the protein surface, the complexes associate to form an array of supercomplex structures, leading to the suggestion that CL acts to steer and “glue” the individual complexes together^30^. However, the precise molecular mechanism responsible for CL-dependent supercomplex stability and activity is unclear.

Monolysocardiolipin (MLCL) is a CL variant that has only three acyl chains and a hydroxyl group in place of the fourth. Accumulation of MLCL is observed in the mitochondrial membranes of people with Barth syndrome (BTHS)^31^. BTHS is caused by a mutation in the tafazzin gene^32^, leading to dysfunctional cardiolipin biosynthesis^33^. Clinical manifestations of BTHS include growth retardation, skeletal muscle fatigue and cardiomyopathy^34,35^. The cellular and molecular basis of BTHS can be attributed to numerous factors, including destabilisation of supercomplexes^36–39^.

*In silico* investigations into the molecular properties of MLCL shows that the lipid retains an equivalent charge state to CL^40^. However, molecular dynamics (MD) simulations reveal contrasting molecular geometries: MLCL exhibits a more cylindrical shape and in the bilayer the phosphate headgroup containing the lyso hydroxyl tilts itself away from the hydrophobic bilayer core and towards the interfacial region. Furthermore, the acyl chain adjacent to the lyso hydroxyl moiety exhibits increased ordering^41^. These molecular properties likely contribute to the observation that MLCL, unlike CL, does not readily localise to negatively curved regions of the membrane^41,42^. Such alterations of the membrane biophysical properties may impact the activity and stability of membrane-embedded proteins^43^.

Early detergent solubilisation studies revealed the reduced affinity of MLCL for CIV^44^. Furthermore, reconstitution of CIV with MLCL retains only 60% of the activity of CL reconstitution. More recently, ^31^P-NMR spectroscopy was used in tandem with detergent solubilisation to reveal MLCL binds with lower affinity to mitochondrial membrane proteins^45^. However, the mechanism by which the unique molecular properties of MLCL impact on specific protein-lipid interactions to affect stability and functionality remains unclear^46^. Moreover, understanding the role of MLCL accumulation in abnormal supercomplex structure and function remains to be understood.

Here, we use CG and atomistic (AT) MD simulations of the *S. cerevisiae* supercomplex CIII2CIV2 (PDB accession code: 6HU9) and murine supercomplex CIII2CIV (PDB accession code: 7O3C) embedded in a mitochondrial membrane mimetic to characterise the interactions of CL and MLCL with the supercomplex. A CG approach is computationally cost-effective and, compared to conventional atomistic simulations, allows for investigation of protein-lipid interactions on longer, more physiologically relevant, timescales. The CG MD simulations showed that CL and MLCL form similar interaction sites on the supercomplex, in particular both lipids form an interaction site at the CIII:CIV interface, also identified in cryo-EM structures. However, free energy calculations suggest that MLCL interactions are weaker than CL. To offer further insight into differing interaction modes between the lipids, atomistic protein-lipid contact analysis was used to probe the contribution of each individual respiratory complex to lipid binding, indicating that MLCL forms a weaker interaction with CIV in the interaction site. The results reveal the distinct protein-lipid interactions of each lipid and may provide an explanation as to why CL may act as a more effective “glue” to stabilise the interface contacts and overall stability of the supercomplex. CG MD of a human CIII2CIV1 supercomplex model demonstrate that the interface site is maintained, bolstering the hypothesis that weaker MLCL interactions at this site could be a mechanism of supercomplex degradation in people with BTHS.

## Methods

### Structures used

Coordinates for *S. cerevisiae* respiratory supercomplex CIII2CIV2 were taken from PDB ID 6HU9^26^. The murine CIII2CIV1 supercomplex was taken from PDB ID 7O3C^29^. The human CIII2CIV1 supercomplex structures was modelled by alignment of human CIII (PDB ID: 5XTE^47^) and CIV (PDB ID: 5Z62^48^) with the murine supercomplex^29^. All bound lipids were removed before processing.

### CG MD simulations

Atomistic protein structures were converted into CG models using MARTINI2.2^49–51^ and the martinize.py script (version 2.4; http://md.chem.rug.nl/index.php/tools/proteins-and-bilayers). An elastic network was applied over the complete supercomplex, between BB beads within 1 nm with a force constant of 1000 kJ mol^-1^ nm^-2^. Protein complexes were embedded in a bilayer of varying lipid ratios and dimensions (Table 1) using insane.py^52^. Standard MARTINI lipid parameters were used for phosphatidyl choline (PC) and phosphatidyl ethanolamine (PE). Tetraoleoyl-CL was taken from ^53^, while trioleoyl-MLCL was taken from reference ^41^.

**Table 1.**
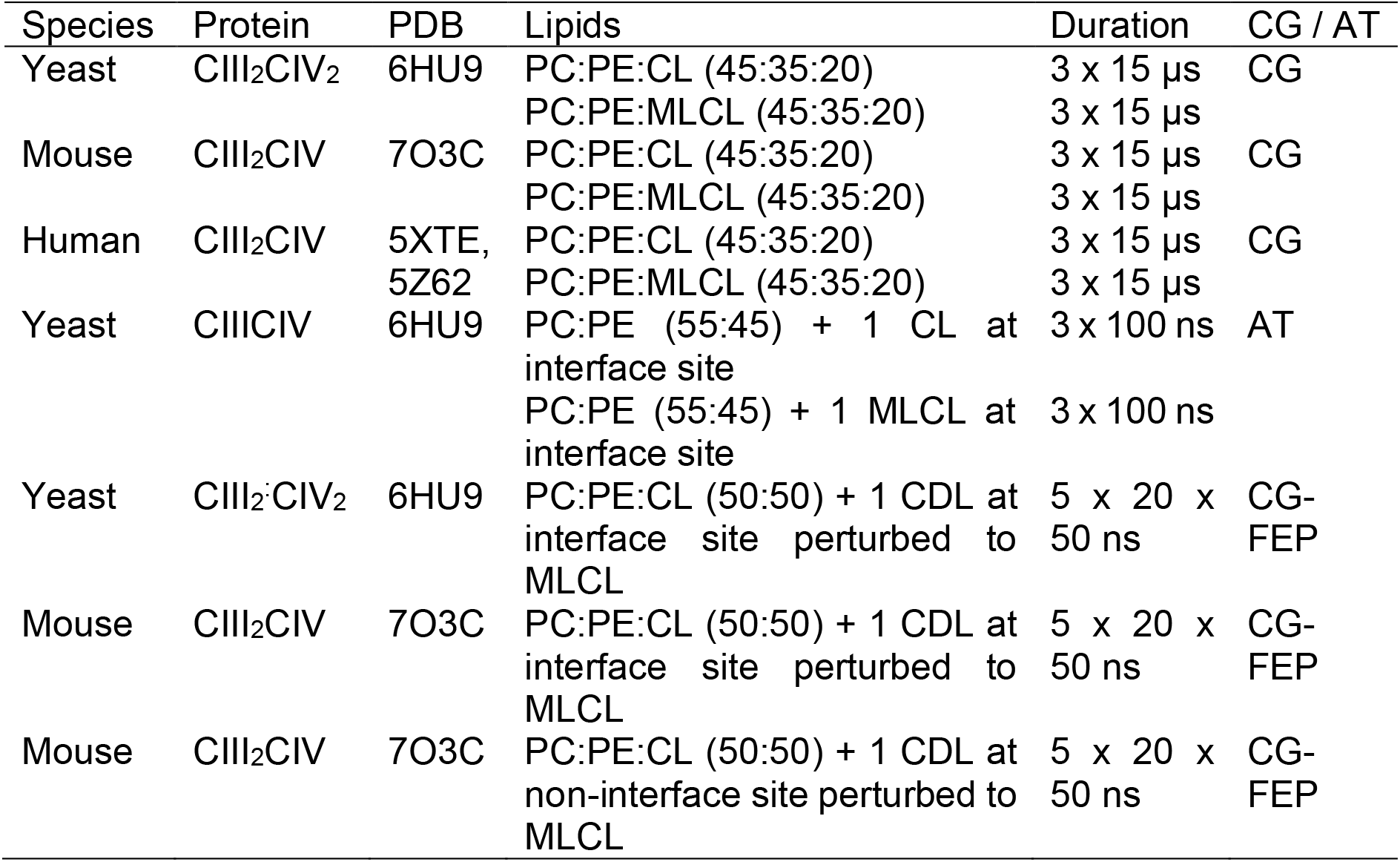
Simulations Performed

| Species | Protein | PDB | Lipids | Duration | CG / AT |
| --- | --- | --- | --- | --- | --- |
| Yeast | CIII<sub>2</sub>CIV<sub>2</sub> | 6HU9 | PC:PE:CL (45:35:20) | 3 x 15 μs | CG |
| | | | PC:PE:MLCL (45:35:20) | 3 x 15 μs | |
| Mouse | CIII<sub>2</sub>CIV | 7O3C | PC:PE:CL (45:35:20) | 3 x 15 μs | CG |
| | | | PC:PE:MLCL (45:35:20) | 3 x 15 μs | |
| Human | CIII<sub>2</sub>CIV | 5XTE, 5Z62 | PC:PE:CL (45:35:20) | 3 x 15 μs | CG |
| | | | PC:PE:MLCL (45:35:20) | 3 x 15 μs | |
| Yeast | CIIICIV | 6HU9 | PC:PE (55:45) + 1 CL at interface site | 3 x 100 ns | AT |
|  |  |  | PC:PE (55:45) + 1 MLCL at interface site | 3 x 100 ns |  |
| Yeast | CIII<sub>2</sub>CIV<sub>2</sub> | 6HU9 | PC:PE:CL (50:50) + 1 CDL at interface site perturbed to MLCL | 5 x 20 x 50 ns | CG-FEP |
| Mouse | CIII<sub>2</sub>CIV | 7O3C | PC:PE:CL (50:50) + 1 CDL at interface site perturbed to MLCL | 5 x 20 x 50 ns | CG-FEP |
| Mouse | CIII<sub>2</sub>CIV | 7O3C | PC:PE:CL (50:50) + 1 CDL at non-interface site perturbed to MLCL | 5 x 20 x 50 ns | CG-FEP |

The CG MD simulations were performed using Gromacs 2020^54,55^. Energy minimisation was performed with the steepest-descent method over 5000 steps. Subsequent two-part equilibration followed. The systems were initially equilibrated for 500 ps with a 10 fs timestep and then for 1000 ps with a 20 fs timestep. Subsequent production simulations (3 x 15 μs for each supercomplex) were run with a 20 fs timestep. For the equilibration, pressure was coupled to a semi-isotropic Berendsen barostat^56^ set to 1 bar, and the temperature was maintained at 323 K via a velocity-rescale thermostat^57^ with a time constant of 1 ps. For the production simulations, pressure was maintained at 1 bar using a Parrinello-Rahman barostat^58^ with a coupling time constant of 12 ps, and a compressibility of 3 x 10^-4^ bar^-1^. The reaction field Coulombic interactions were switched off at 1.1 nm. Van der Waals interactions were calculated using a 1.2 nm cut-off with a switching function in place from 0.9 nm. LINCS constraints were applied to covalent bonds.

### Free energy perturbation (FEP) calculations

Free energy perturbation (FEP) calculations were performed on the *S. cerevisiae* supercomplex structure with a single CIII monomer and its neighbouring CIV monomer, or on the full murine supercomplex. The starting structure was obtained from a snapshot of a 15 μs simulation where a single CL was bound at the interface position, or at the equivalent non-interface site for the murine supercomplex.

The CL was alchemically perturbed into a MLCL molecule by removing one of the four acyl chains, see Supplementary data. FEPs were run as previously described ^59^. In brief, 21 windows were used to perturb the Lennard-Jones interactions in even sized steps of 0.05. Each window was energy minimised using the steepest descent method and then simulated for 50 ns using the same conditions as for the unbiased simulations. Softcore potentials were applied to the Lennard-Jones interactions using an α of 0.5 and a σ of 0.3. The results were analysed using the *alchemical analysis* script and implemented a Multistate Bennett acceptance ratio (MBAR) method to calculate the free energy landscape^60^. To obtain a *ΔΔG_binding_* value, alchemical transformation was performed both in the binding site (calculating *ΔG_bound_*) as well as for a lipid in a membrane with no protein present (calculating *ΔG_free_*). This process was repeated five times for both bound and free simulations.

### Atomistic MD simulations

The *S. cerevisiae* CIII1CIV1 supercomplex structure with a single CL bound at the CIII:CIV interface was back-mapped from a CG structure into atomistic detail using CG2AT2^61^, with adaptation to include the MLCL topology^41^. The initial CG coordinates were obtained from selecting the final frame of the 15 μs CIII1CIV1 supercomplex simulation trajectory. This was followed by a steepest descent energy minimisation for 50000 steps. Simulations were performed with GROMACS 2020^54,55^ using the CHARMM36m forcefield^62^. Particle Mesh Ewald^63^ was used for modelling long-range electrostatics. Pressures were maintained at 1 bar using Parrinello-Rahman barostat^58^ with a coupling constant of 2 ps and temperatures at 310 K using a velocity-rescale thermostat^57^ and a coupling constant of 0.2 ps. Short range van der Waals and electrostatics were cut-off at 1.2 nm and bonds were constrained using the LINCS algorithm^64^.

### Analysis

Analysis was performed using inbuilt tools provided by GROMACS 2020^54,55^, VMD^65^, and MDAnalysis^66,67^. The VMD plugin VolMap was used for initial assessment of lipid interaction sites, using a resolution of 2 Å for the grid and particle size 3x the particle radius. Densities were calculated for all frames and combined using the mean. Lipid binding sites were identified and the kinetics of dissociation were assessed using PyLipID^68^. Visualisation and image generation was carried out using VMD^65^ and PyMOL.

## Results

To investigate CL and MLCL interaction with mitochondrial supercomplexes, 3 x 15 μs CG MD simulations were run of yeast and mammalian CIII_2_CIV_1-2_ supercomplex structures, in bilayers containing PC, PE and either CL or MLCL.

### CL and MLCL interact at sites identified experimentally

Volumetric maps assessing the average weighted density of each lipid’s phosphate beads reveal defined lipid binding sites for both CL (Figure 1B) and MLCL (Figure S1). Areas of lipid density are observed in the inner cavity of CIII, which is known to be highly hydrophobic, and contains PC, PE and CL in cryo-EM structures (Figure 1A). The areas of lipid occupancy that do not correspond to bound CL in the cryo-EM structure are generally located on the protein surface exposed to the membrane, and therefore are more susceptible to removal during protein purification procedures.

### CL and MLCL interact at the CIII:CIV interface

Lipid density is observed at the interface region between the complex III and IV (Figure 1B), in the same region that a bound CL lipid was identified in the cryo-EM yeast and murine structures (Figure 1A).

To gain more detailed insights into this site, the CG data were analysed using the PyLipID program, which groups protein residues into binding sites by clustering residues that frequently interact simultaneously with the same lipid molecule^68^. For the yeast supercomplex, PyLipID identifies a number of sites including a very high occupancy site at the interface region. This site contains three lysine residues from CIII (Rieske subunit K44, K51; Subunit 8 K35), a lysine from CIV (COX5A K97) and several polar residues from both CIII and CIV (Figure 1B, Table 2), which interact with the CL headgroup (ie. phosphate groups and bridging glycerol). For the murine supercomplex, PyLipID also identifies a high occupancy site at this interface region, which contains an arginine from CIII (Subunit 8 R24), two arginines and a lysine from CIV (COX3 R156, COX6A2 R17, COX7A2L K56), and several polar residues from both.

**Table 2.** Residues identified in the interface interaction site as identified by PyLipID. All residues with occupancy of >40% are listed. Note that the yeast supercomplex, due to having two CIV monomers, also has two interface interaction sites.

| Species | Lipid | Residues |
| --- | --- | --- |
| Yeast | CL | Site 1: CIII: Rieske subunit: K44, K51; Sub8: K35; Sub9: S2, F3, S4, S5<br>CIV: COX5A: S93, F94, K97 |
|  |  | Site 2: CIII: Rieske subunit: K44, K51; Sub8: K35; Sub9: S2, S4, S5<br>CIV: COX5A: S93, K97 |
|  | MLCL | Site 1: CIII: Rieske subunit: K44, N46, K51; Sub8: K35<br>cCIV: COX1: K408; COX5A: G90, S93, K97 |
|  |  | Site 2: CIII: Rieske subunit: K44, N46, K51, Y55; Sub8: K35, P36, L37, Q38, H42<br>CIV: COX5A: G90, S92, S93, F94, K97 |
| Mouse | CL | CIII: Sub8: R24, F26, S28, S31<br>CIV: COX7A2L: S54, G55, K56; COX3: R156, N157, H158, Q161, L223; COX6: S3; COX6A2: R17 |
|  | MLCL | CIII: Sub8: R24, S31<br>CIV: COX7A2L: Y53, S54, G55, K56; COX3: R156, N157, H158, N160, L223; COX6: S3 |
| Human | CL | CIII: Rieske subunit: R93, F113; Sub4: T305, H309; Sub8: R25, A26, Y27, P28, H29, V30, T32<br>CIV: COX3: N157, Q158; COX5B: A32; COX6A1: R36, K39 |
|  | MLCL | CIII: Rieske subunit: R93, F113; Sub4: T305, H309; Sub8: R25, A26, Y27, P28, H29, V30, T32<br>CIV: COX3: N157, Q158; COX5B: A32; COX6A1: R36, K39 |

In simulations where CL was replaced by MLCL (Figure 2A), analysis using both volumetric maps and PyLipID also identified the interface interaction site as being a site for the MLCL headgroup (Figure 2B, Figure S1, Table 2), indicating that both CL and MLCL are able to bind the interface site.

**Figure 2.**
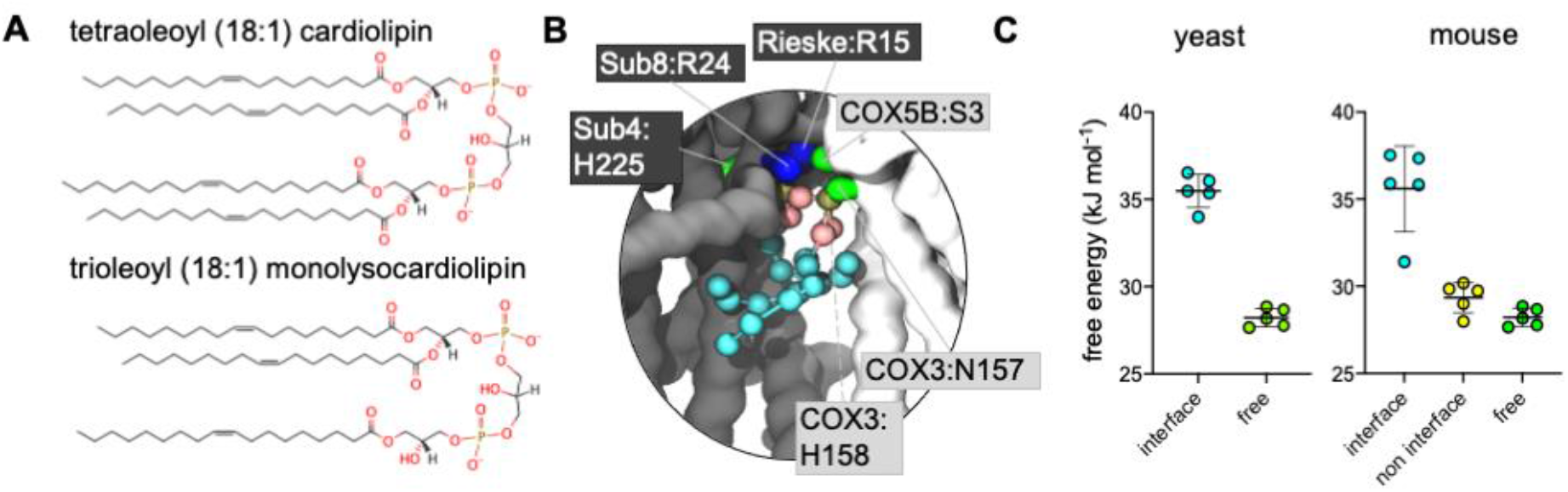
**A)** Structures of CL and MLCL used for this study. **B)** CG MLCL pose used for free energy perturbation (FEP) demonstrating MLCL interaction in the murine supercomplex interface. Protein residues within 5.2 Å of MLCL headgroup (phosphate and bridging glycerol beads) have sidechains shown. **C)** CG FEP of CL vs MLCL in yeast and mouse (the latter has equivalent interface and non-interface sites)

### CL interacts more favourably than MLCL at the interface site

Given that the presence of MLCL in place of CL is thought to destabilise these supercomplexes, we reasoned that the site identified on the interface between CIII and CIV might be of physiological relevance. Therefore, free energy perturbation (FEP) calculations were used to characterise the binding energies of CL and MLCL to the protein interface site.

To identify the relative difference in binding affinity of CL and MLCL for the interface site, FEPs were performed in which CL was alchemically transformed into MLCL (Figure S2) both bound to the protein and free in the membrane. The *ΔG* associated with transformation of protein-bound CL into MLCL was higher than the *ΔG* for transformation in the bulk bilayer, suggesting that CL interacts with the protein interface site more favourably than MLCL, with a *ΔΔG* value calculated at 8 kJ mol^-1^ (Figure 2C). This is roughly equivalent to the binding strength of CL to certain other proteins ^69^, suggesting that it is a substantial difference.

### Presence of CIV determines preference for CL at interface site

The murine supercomplex contains only one copy of CIV, but two copies of CIII (an obligate dimer). Thus, there will be one CIII-CIV interface site, but also a site in the equivalent position on the other CIII monomer, where no neighbouring CIV is present, which we term the ‘non-interface site’. PyLipID identifies the non-interface site as a lipid binding site. However FEP calculations show that at the non-interface site the preference for CL over MLCL is no longer apparent (Figure 2C). This suggests that CIV is important in providing the preference for CL *vs.* MLCL at the interface site.

### CL makes more contacts than MLCL with CIV at the interface site

In order to further understand the importance of CIV for the interface site, atomistic simulations of the CIII:CIV interface with a single CL or MLCL molecule were performed. Snapshots from the simulations indicate that the four acyl chains of CL may be able to form more extensive contacts with CIV than is possible for MLCL with only three acyl chains (Figure 3A, B). Contact analysis shows that the number of CIII-lipid contacts are similar for CL and MLCL (Figure 3C). However, MLCL makes fewer contacts with CIV than CIII. This suggests that the preference for CL vs MLCL in the interface site is driven by a greater number of CIV-acyl chain interactions for CL as compared to MLCL, which supports the conclusions from the free energy analyses.

**Figure 3.**
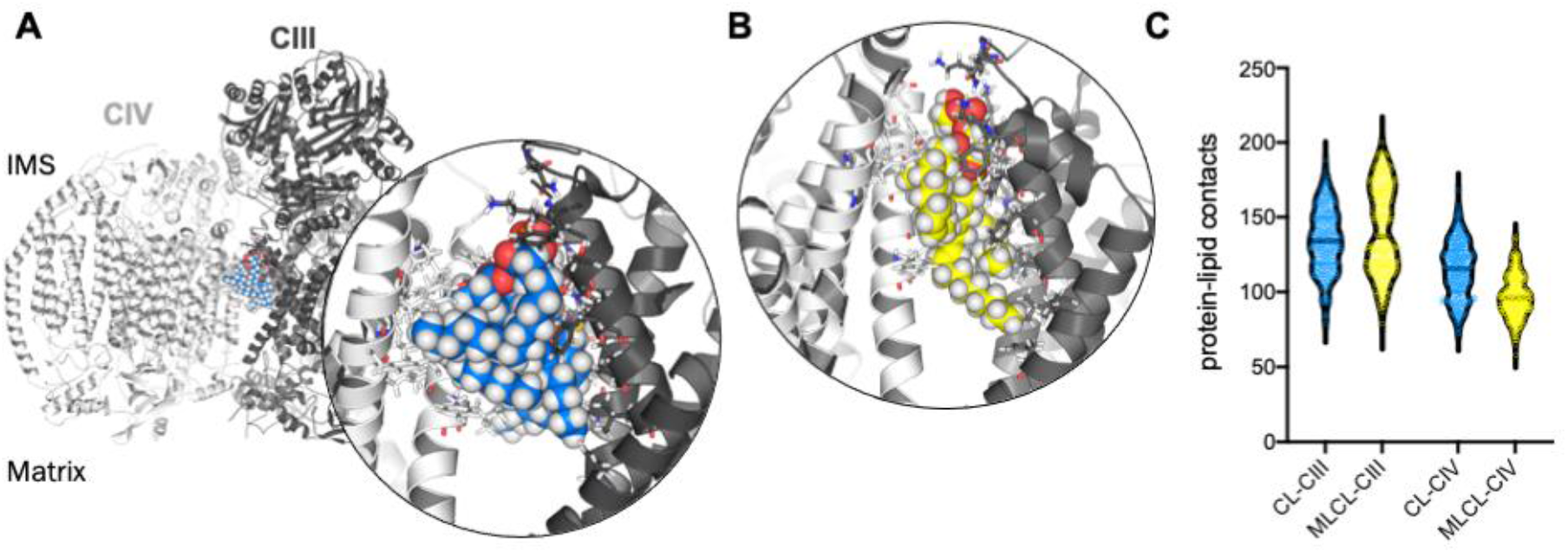
**A)** Atomistic yeast supercomplex with interface cardiolipin shown in blue. Inset, all protein residues within 4 Å of the CL are shown as sticks. **B)** Atomistic simulation of the yeast supercomplex with MLCL, shown in yellow. MLCL makes fewer contacts with CIV (white) than CIII (grey). **C)** Number of contacts made in atomistic simulations between CIII or CIV and either CL or MLCL. CL and MLCL make similar contacts to CIII (median atoms in contact are 134 and 137), whereas MLCL makes substantially fewer contacts to CIV (median atoms in contact are 115.5 and 96).

### Interface site identified in a human model of CIII2CIV1

In order to confirm that the lipid interaction site at the CIII:CIV interface could indeed be relevant for BTHS, we ran CG MD simulations of a model of the human CIII2CIV1 supercomplex. Whilst a structure of human respirasome has been resolved (ie containing CI as well as CIII and CIV) no structure of the CIII2CIV1 exists to our knowledge. Therefore, we built a model based on human CIII and CIV structures^47,48^ superposed on the murine supercomplex^29^. Analysis using VolMap (Figure 4A) and PyLipID (Figure 4B) again identified the interface interaction site. As for the yeast and murine complexes, the human interface interaction site contains two arginine residues from CIII and a further positively charged residue on CIV (Figure 4B and Table 2). Identification of the interface lipid interaction site suggests that the interaction of CL and MLCL in the human supercomplex may be similar to that seen in yeast and murine supercomplexes.

**Figure 4.**
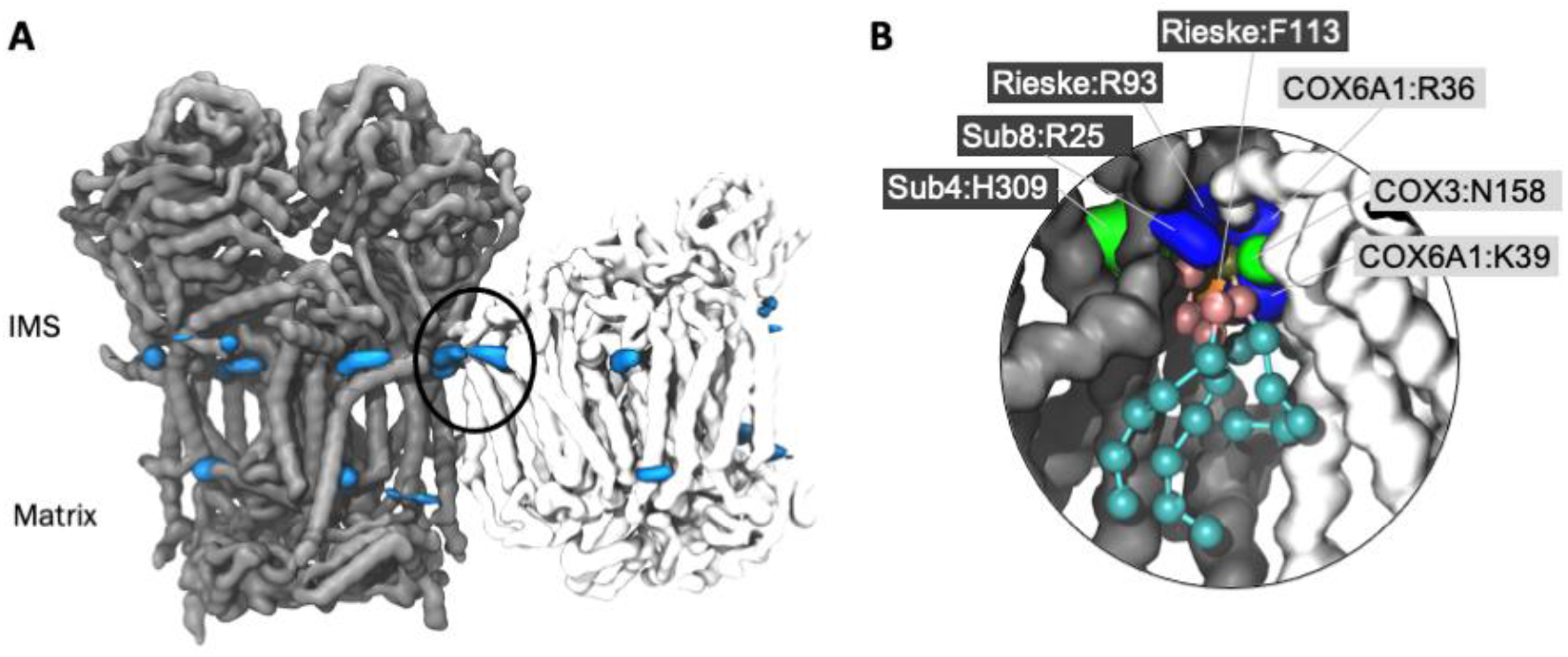
**A)** CL headgroup density from human CIII2CIV CG MD simulations. CIII backbone is shown in grey and CIV in white, with CL headgroup densities in blue. **B)** Snapshot of CL in the human supercomplex interface interaction site (location circled in A). Protein residues within 5.2 Å of CL headgroup have sidechains shown. CL is shown with phosphate moieties in tan, glycerol moieties in pink, and acyl chain beads in cyan. The backbone is shown of CIII (grey) and CIV (white), with sidechains shown of residues within 5.2 Å of the CL headgroup; lysine and arginine sidechains in blue, histidine and asparagine in green, and phenylalanine in orange.

## Discussion

### CL and MLCL association with the mitochondrial supercomplex CIII_2_CIV_1-2_

The association of CL with respiratory complexes and supercomplexes is well-documented^70^. MLCL has been shown to interact with CIV but with a lower affinity than CL^44^ and reconstitution of both CIV and the ATP/ADP transporter with MLCL rather than CL lead to reduced protein activity^44,71^. However, there has been less investigation into the protein-lipid interactions of supercomplexes and MLCL. Here, we have shown that both CL and MLCL show enrichment at defined binding sites, consistent with sites identified previously in cryo-EM structures and MD simulations focussed on the individual respiratory complexes^72,73^.

Both CL and MLCL appear to show greater enrichment in the matrix-facing leaflet compared to the inter membrane space (IMS)-facing leaflet. This is likely due to the physiological distribution of CL in the inner mitochondrial membrane in which it represents up to 20% of the matrix-facing leaflet and presents substantial paucity in the IMS-facing leaflet^7^, an observation which is also apparent in bacterial proteins^74^. Therefore, it is plausible that functionally significant CL binding would have evolved to occur in the matrix-facing side of the supercomplex protein.

### CL and MLCL interaction at the CIII:CIV interface site

The interface site is formed of a cluster of positively charged residues, alongside several polar residues. The CL glycerol moieties that connect phosphate groups with acyl tails also interact with aromatic groups such as yeast W286 on subunit 4 of CIII, and murine F35 on the Rieske subunit of CIII (see for comparison Fig 1A). Together these interactions form all the characteristic features of bacterial CL interaction sites as we previously identified^74^ suggesting a conserved mode of binding.

The CL interface site is conserved across yeast, mouse and human CIIII_2_:CIV_1-2_ supercomplexes (Figures 1, 4 and S1) indicating functional importance. The site is also conserved for MLCL. This is likely because the site is ideally suited for the MLCL headgroup, which is identical in chemical composition to that of CL. However, as we show, MLCL does overall have a weaker interaction at this site than CL.

It has been suggested that lipid enrichment at interface sites between complexes may be responsible for “gluing” the complexes together^20^. FEP calculations of the interface site vs. the equivalent non-interface site on the other CIII monomer (Figure 3C) suggest that the presence of CIV contributes to the relative strength of association of CL interaction *vs.* that of MLCL. Analysis of atomistic data also suggests that, compared with MLCL, CL forms more contacts with CIV, but no difference in the number of contact with CIII. Residues interacting with the lipid headgroups do not differ greatly for CL and MLCL, thus the difference in interactions may be attributed to the extra acyl chain of CL, which leads to CL exhibiting a less conical shape than MLCL. CL may therefore be able to maximise its stability by bridging the two protein surfaces.

It is plausible that this lack of defined bridging interactions across the interface causes MLCL to not only bind more weakly to the supercomplex surface, but may also influence the supercomplex stability. Indeed, CL has previously been suggested to act as a supercomplex glue^20,72^ as well as extend the minimal protein-protein interaction surface area^26^. For this mechanism to be effective, the lipid would be required to bridge both complexes stably, and the acyl chains of MLCL do not exhibit this behaviour. FEP calculations also suggest CL binds more strongly to the interface site. This has implications for the supercomplex stability in general; our finding that CL has a higher affinity for this site than MLCL suggests that CL is likely to spend more time at the interface site, therefore provide more stability to the supercomplex over long timescales.

Interestingly, previous PMF calculations indicate CL and MLCL exhibit identical free energies of binding to a CIV binding site proposed to be involved in supercomplex interface oligomerisation^30^. The discrepancy with our data could be accounted for if one considers our results that suggest the interface binding site requires the presence of both complexes to stably bind CL over MLCL.

MD simulations show CL enrichment at interface sites on individual respiratory complexes prior to oligomerisation into supercomplex structures^30^. Furthermore, the site on CIII has been identified in previous simulations of only the CIII dimer (site III in ref. ^72^), and the CIV component of the interface site has also been observed in simulations of CIV alone (site V in ref. ^73^), perhaps indicating that CL interaction precedes supercomplex formation. However, recent work has shown that supercomplex assembly may happen in tandem with assembly of the individual complexes^75^, rather than proceeding after assembly of constituent components. Thus, it is unclear whether CL helps to guide supercomplex assembly, or functions principally to stabilise existing supercomplexes.

### Limitations

This work does not account for CL in all supercomplex variants; other supercomplexes (in particular the mammalian respirasome, containing CI as well as CIII2 and CIV^47,76–79^) have a different CIII-CIV interface: where CIII interacts with CIV in CIII_2_CIV_1-2_, instead CIII instead forms an interaction with CI. Furthermore, in the recently structurally resolved CICIICIII2CIV2 ciliate supercomplex, there is no CIII-CIV interface at all ^80^. Thus, the interface lipid sites will be different, although the same mechanism for weaker MLCL interaction is conceivable.

In this study, we show that MLCL interacts less favourably than CL in the supercomplex interface interaction site. Although we do not demonstrate how CL or MLCL contribute to the free energy of interaction of CIII with CIV, our results support the hypothesis that CL and not MLCL glue the respiratory complexes together^20,21,23^, and are in agreement with data showing that MLCL does not bind as tightly to mitochondrial proteins^45^.

## Conclusions

In summary, our results show that CL, with its conical shape, can bridge the CIII:CIV interface in CIII_2_CIV_1-2_ more successfully than the more cylindrical MLCL, thereby stabilising these supercomplexes. Our results, for the first time, provide molecular understanding to the lack of supercomplexes in BTHS, where MLCL accumulates and mature CL is depleted. While the role of supercomplexes remains to be fully understood, the results presented here demonstrate the importance of specific lipid interactions in supercomplexes, and pinpoint how a relatively minor change to a lipid species might have system-wide implications, such as in BTHS.

## Author contributions

Author contributions, described using the CASRAI CRedIT typology (http://casrai.org/credit), are as follows: conceptualisation: A.L.D.; validation: R.A.C., N.H.; formal analysis: R.A.C., N.H.; investigation: R.A.C., N.H.; provision of computing resources: P.J.S., M.S.P.S.; data curation: R.A.C., A.L.D.; writing – original draft preparation: R.A.C., N.H., A.L.D.; writing – review & editing: R.A.C., N.H., P.J.S., M.S.P.S., A.L.D.; visualization: R.A.C., N.H., A.L.D.; supervision: R.A.C., A.L.D.; project administration: A.L.D.; funding acquisition: P.J.S., M.S.P.S..

## Acknowledgements

R.A.C., A.L.D., P.J.S., and M.S.P.S. are funded by Wellcome (208361/Z/17/Z). A.L.D. was also supported by the Department of Biochemistry, University of Oxford. Research in the M.S.P.S.’s group is also supported by BBSRC and EPSRC. Research in P.J.S.’s laboratory is funded by the MRC (MR/S009213/1) and BBSRC (BB/P01948X/1, BB/R002517/1, and BB/S003339/1). Simulations were carried out, in part, on ARCHER, provided by HECBioSim, the UK High End Computing Consortium for Biomolecular Simulation (hecbiosim.ac.uk), which is supported by the EPSRC (EP/L000253/1). We would like to thank Dr. Irfan Alibay and Dr. Michael Horrell for the maintenance of local computing resources.

