## Supplementary Information for "Cardiolipin, and not monolysocardiolipin, preferentially binds to the interface of Complexes III and IV"

### Supplemetary Figures

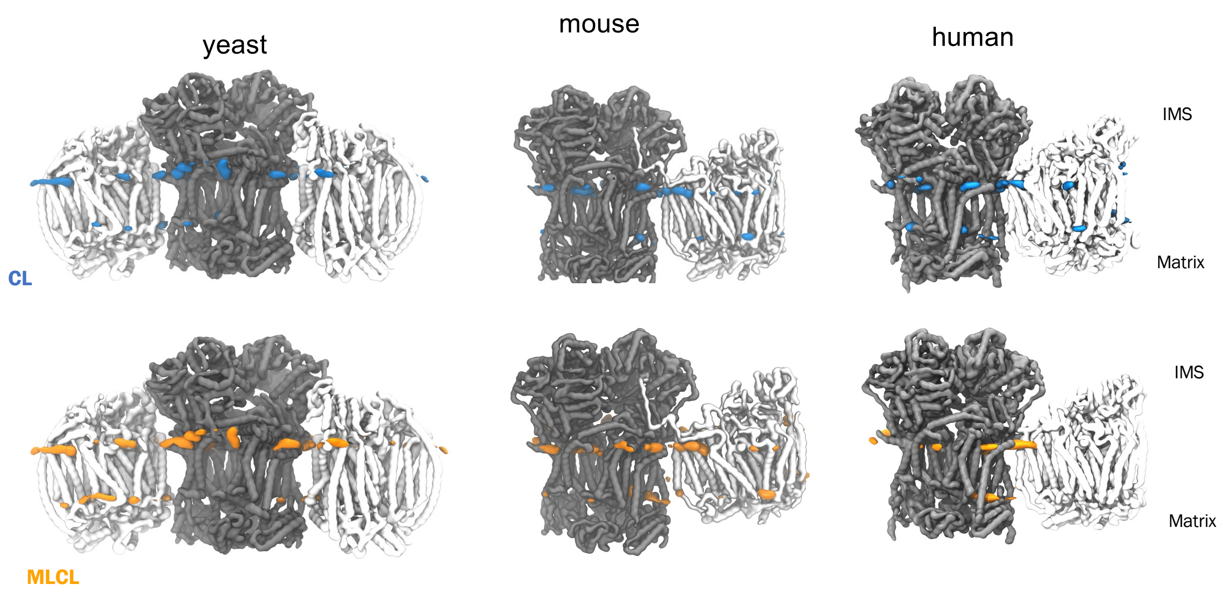


**Figure S1.** CL (top row) and MLCL (second row) headgroup density from yeast, mouse and human $CIII_2CIV_{1\text{-}2}$ CG MD simulations. CIII backbone is shown in grey and CIV in white, with CL headgroup densities in blue and MLCL densities in orange.

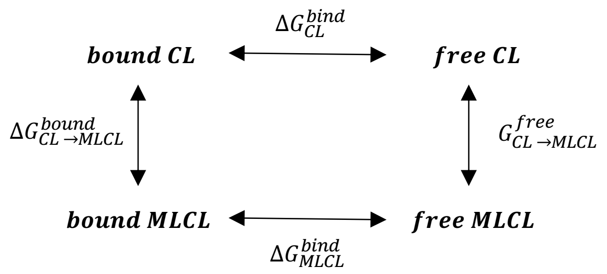


$$\Delta\Delta G = \Delta G_{CL}^{bind} - \Delta G_{MLCL}^{bind}$$
$$= \Delta G_{CL\rightarrow MLCL}^{bound} - \Delta G_{CL\rightarrow MLCL}^{free}$$

**Figure S2.** Thermodynamic cycle used for the free energy perturbation simulations.

**Supplementary Data:**

1. Coordinates of CL and MLCL binding sites, as identified by PyLipID, for yeast, mouse and human supercomplexes (6 files)

2. Itp file for CL > MLCL used for FEP
